# JAM-A functions as a female microglial tumor suppressor in glioblastoma

**DOI:** 10.1101/761445

**Authors:** Soumya M. Turaga, Daniel J. Silver, Defne Bayik, Evi Paouri, Sen Peng, Nozha Borjini, Sarah Stanko, Ulhas Naik, Ruth A. Keri, James R. Connor, Jill S. Barnholtz-Sloan, Joshua B. Rubin, Michael Berens, Dimitrios Davalos, Justin D. Lathia

**Affiliations:** Department of Biological, Geological, and Environmental Sciences, Cleveland State University, Cleveland, Ohio, USA; Department of Cardiovascular & Metabolic Sciences, Lerner Research Institute, Cleveland Clinic Cleveland, Ohio, USA; Case Comprehensive Cancer Center, Case Western Reserve University, Cleveland, Ohio, USA; Department of Neurosciences, Lerner Research Institute, Cleveland Clinic Cleveland, Ohio, USA; Cancer and Cell Biology Division, TGen, Phoenix, AZ 85004, USA; Cardeza Center for Vascular Biology, Department of Medicine, Thomas Jefferson University, Philadelphia, PA, 19107, USA; Departments of Pharmacology and Genetics and Genome Sciences, Case Western Reserve University, Cleveland, Ohio; Department of Neurosurgery, Penn State College of Medicine, Hershey, PA, United States; Department of Population and Quantitative Health Sciences, Case Western Reserve University School of Medicine, Cleveland, Ohio; Department of Pediatrics, Washington University School of Medicine, St. Louis, MO 63110, USA; Department of Molecular Medicine, Cleveland Clinic Lerner College of Medicine of Case, Western Reserve University, Cleveland, Ohio, USA; Rose Ella Burkhardt Brain Tumor and Neuro-Oncology Center, Cleveland Clinic, Ohio, USA

**Author notes:** **Correspondence:** Dr. Justin D. Lathia, Lerner Research Institute, 9500 Euclid Ave, NC10, Cleveland, OH 44195, USA.

**Keywords:** glioblastoma, microglia, junctional adhesion molecule-A, sex differences

## Abstract

Glioblastoma (GBM) remains refractory to treatment. In addition to its cellular and molecular heterogeneity, epidemiological studies indicate the presence of additional complexity associated with biological sex. GBM is more prevalent and aggressive in male compared to female patients, suggesting the existence of sex-specific growth, invasion, and therapeutic resistance mechanisms. While sex-specific molecular mechanisms have been reported at a tumor cell-intrinsic level, sex-specific differences in the tumor microenvironment have not been investigated. Using transgenic mouse models, we demonstrate that deficiency of junctional adhesion molecule-A (JAM-A) in female mice enhances microglia activation, GBM cell proliferation, and tumor growth. Mechanistically, JAM-A suppresses anti-inflammatory/pro-tumorigenic gene activation via interferon-activated gene 202b (Ifi202b) and found in inflammatory zone (Fizz1) in female microglia. Our findings suggest that cell adhesion mechanisms function to suppress pathogenic microglial activation in the female tumor microenvironment, which highlights an emerging role for sex differences in the GBM microenvironment and suggests that sex differences extend beyond previously reported tumor cell intrinsic differences.

**Summary:** Turaga et al. demonstrate that female microglia drive a more aggressive glioblastoma phenotype in the context of JAM-A deficiency. These findings highlight a sex-specific role for JAM-A and represent the first evidence of sexual dimorphism in the glioblastoma microenvironment.

## Introduction

Glioblastoma (GBM) is the most common primary malignant brain tumor and, despite an aggressive standard of care, has a median survival between 15 and 20 months (Stupp et al.,2017). There are multiple barriers to the development of more effective therapies, including inter- and intra-tumoral heterogeneity at the cellular and molecular levels, a high degree of invasion into the surrounding brain parenchyma, mechanisms of intrinsic resistance to radiation and chemotherapy, and an immune-suppressive tumor microenvironment. The GBM microenvironment consists of 30-50% microglia, the resident immune cells of the brain, and infiltrating tumor-associated macrophages (TAMs) (Roesch et al., 2018b). The interaction between GBM cells and microglia/TAMs is principally mediated through direct cell-cell contact and a series of secreted factors, with GBM cells amplifying the immune-suppressive phenotypes of microglia/TAMs and microglia/TAMs concomitantly driving GBM cell growth and tissue infiltration (Arcuri et al., 2017; Guadagno and Presta, 2018; Sorensen and Dahlrot, 2018). An understudied barrier to effective treatment is the inherent sex differences that exist within GBM (Ippolito et al., 2017;Kfoury et al., 2018; Sun et al., 2015; Sun et al., 2014). These differences are supported at the epidemiological level, with the male to female incidence ratio being 1.6:1 (Gittleman et al., 2017; Ostrom et al., 2016; Sun et al., 2015). Furthermore, these differences manifest clinically, with females showing a more dispersive phenotype radiographically (Yang et al., 2019) and males experiencing a poorer prognosis (Ostrom et al., 2018). There is supporting evidence in the literature suggesting that sexual dimorphism in GBM is mediated through sex-specific differences in tumor cell-intrinsic oncogenic signaling pathways (Kfoury et al., 2018;Ostrom et al., 2019; Ostrom and Kinnersley, 2018; Sun et al., 2014), epigenetic states, and metabolic profiles (Ippolito et al., 2017). While sex differences might induce differential GBM cell-intrinsic responses, leading to differences in survival, sex-specific differences in the tumor microenvironment have not yet been elucidated.

Microglia are a major cell population in the tumor microenvironment with inherent sex signatures that impact their development, maintenance, activation and overall function in homeostatic and disease states (Lenz and McCarthy, 2015). Microglia-mediated sex differences are more prominent and have been well characterized in neurological disorders such as Autism and Alzheimer’s disease (Hanamsagar and Bilbo, 2016). Recent studies have confirmed that microglia continuously survey their surrounding environment (Davalos et al., 2005) through a variety of cell adhesion mechanisms (Meller et al., 2017), and this extends to interactions with adjacent tumor cells (Silver et al., 2016). These interactions are mediated in part by tight junction proteins, among which junctional adhesion molecule-A (JAM-A, also known as F11r) was shown to be highly expressed by tumor-associated microglia and TAMs (Pong et al., 2013). We previously demonstrated that JAM-A was necessary and sufficient for GBM cancer stem cell maintenance (Alvarado et al., 2016; Lathia et al., 2014), but its role in the tumor microenvironment has not been examined. Here, we specifically investigated whether JAM-A could mediate sex differences in tumor progression, and found that JAM-A deficiency in microglia impacts GBM pathogenesis in a sex-specific manner.

## Results and Discussion

### JAM-A deficient female mice display an aggressive GBM phenotype

To assess the function of JAM-A in the tumor microenvironment, we took advantage of JAM-A deficient mice in combination with a transplantable syngeneic orthotopic mouse glioma model. We transplanted an equivalent number of GL261 cells into 6-week-old male and female wild-type and JAM-A deficient mice and assessed survival based on the development of neurological signs, which reflected the experimental endpoint (**Fig. 1A**). Using this approach, no differences were observed in the survival of wild-type and JAM-A deficient mice when both sexes were combined together (**Fig. 1B**). Given the sex differences in survival observed in human patients, we compared the groups based on sex of the mice. When comparing the wild-type groups, we observed a similar trend with GL261 cells as seen in human population, where females survived significantly longer compared to males (**Fig. 1C**). In the context of JAM-A deficiency, however, disease aggression was significantly increased in females compared to males (**Fig. 1D**), reversing the survival difference observed between male and female GBM patients. When comparing genotypes by sex, JAM-A deficient females had significantly poorer survival compared to wild-type females in both the GL261 and SB28 models (**Fig. 1E, Supplemental Fig. 1A**). However, this difference was not observed between JAM-A deficient and wild-type males (**Fig. 1F**). Importantly, a single copy of JAM-A was sufficient to improve the survival of female mice, as the heterozygous JAM-A mice had a significantly better survival than JAM-A deficient mice (**Supplemental Fig. 1B**). As expected, this was not observed in the context of male heterozygous and JAM-A deficient mice (**Supplemental Fig. 1C**). Taken together, these data suggest a tumor-suppressive role for JAM-A specifically in the female tumor microenvironment.

**Figure 1:**
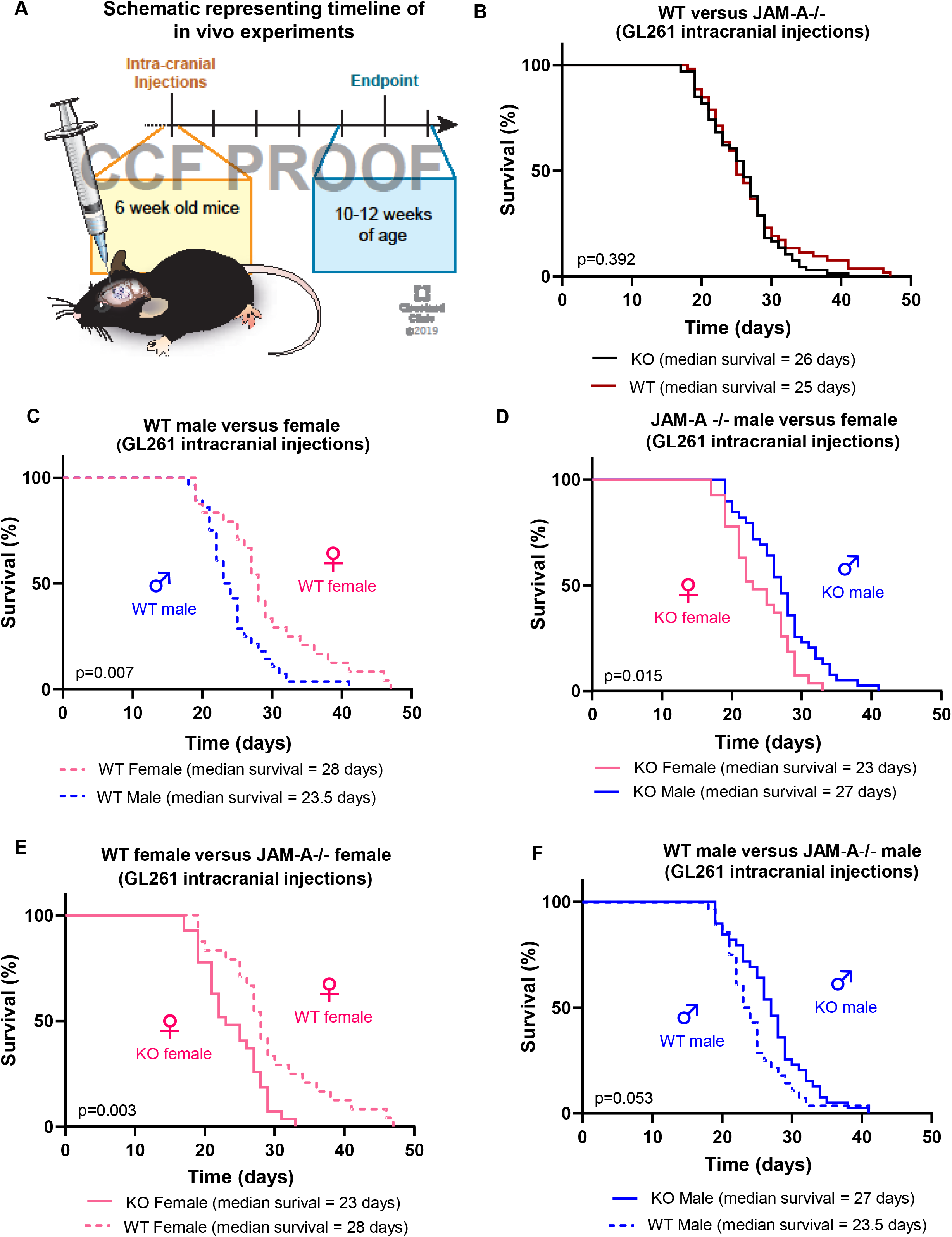
Female JAM-A deficient mice exhibit poor survival upon tumor implantation. A) Schematic showing the timeline of intracranial implantation of tumor cells and manifestation of end-point symptoms. B) Survival curves upon intracranial implantation of GL261 cells into wild-type (n=52, median survival=25 days) and JAM-A deficient mice (n=66, median survival=26 days); *p=0.392*. C) Survival curves upon intracranial implantation of GL261 glioma cells into wild-type male (n=28, median survival=23.5 days) and wild-type female mice (n=24, median survival=28 days); *\*\*p=0.007*. D) Survival curves upon intracranial implantation of GL261 glioma cells into JAM-A deficient male (n=39, median survival=27 days) and JAM-A deficient female mice (n=27, median survival=23 days); *\*p=0.015*. E) Survival curves of JAM-A deficient (n=27, median survival=23 days) and wild-type females (n=24, median survival=28 days); *\*\*p=0.003*. F) Survival curves between JAM-A deficient (n=39, median survival=27 days) and wild-type male mice (n=28, median survival=23.5 days); *p=0.053*. Kaplan-Meier survival curves were plotted using GraphPad Prism 6.0, and the p-value was assessed using log-rank (Mantel-Cox) test.

### Female JAM-A deficient mice have increased activation of microglia in the tumor microenvironment compared to other genotypes

These sex differences in survival of mice implanted with identical cells suggest that the host sex and microenvironment are likely to be the mediators of the observed effect. We next assessed the potential for cells in the tumor microenvironment to drive these differences. We first focused on microglia given their reported sex differences in developmental, homeostatic, and disease states, including optic pathway glioma (Toonen et al., 2017a), and we and others have previously reported that GBM-associated microglia express JAM-A (Lathia et al., 2014; Pong et al., 2013). To determine the expression of JAM-A in various cell populations, we assessed the Brain RNA-seq database that provides expression levels for purified cell types from the healthy adult mouse brain (Zhang et al., 2014; Zhang et al., 2016). This analysis revealed that JAM-A expression was highest in microglia, followed by endothelial cells in the brain in both mouse and humans (**Fig. 2A**), consistent with the previous report of JAM-A expression in various types of endothelial cells (Naik et al., 2003). To determine whether there were baseline differences in microglial number in adult wild-type and JAM-A deficient mice, we assessed the number of microglia via immunofluorescence for Ionized calcium binding adaptor molecule 1 (Iba1) positive cells. We found that JAM-A deficient mice had significantly more Iba1+ cells compared to wild-type controls, but there was no difference between males and females (**Fig. 2B**). To determine whether there were differences in activation state, we assessed adult wild-type and JAM-A deficient mice transplanted with GBM cells and found that female JAM-A deficient mice had a significant increase in amoeboid/rounded Iba1+ cells, a surrogate of microglial activation (Jonas et al., 2012), compared to all other genotypes (**Fig. 2C, D**). JAM-A deficient female microglia also showed higher expression of the microglia activation marker found in inflammatory zone (Fizz1) at baseline (**Fig. 2E**). Fizz1 expression in microglia and TAMs correlates with an anti-inflammatory and pro-tumorigenic state in the tumor microenvironment (Grimaldi et al., 2019; Roesch et al.,2018a; Zhou et al., 2015). We also assessed JAM-A deficient and wild-type female mice for other immune cells by flow cytometry and did not observe differences in immune populations in the blood and brain (**Supplemental Fig. 2**). These data indicate that JAM-A deficiency results in enhanced microglia activation specifically in JAM-A deficient female mice, which experience aggressive GBM growth and poor survival compared to other genotypes.

**Figure 2:**
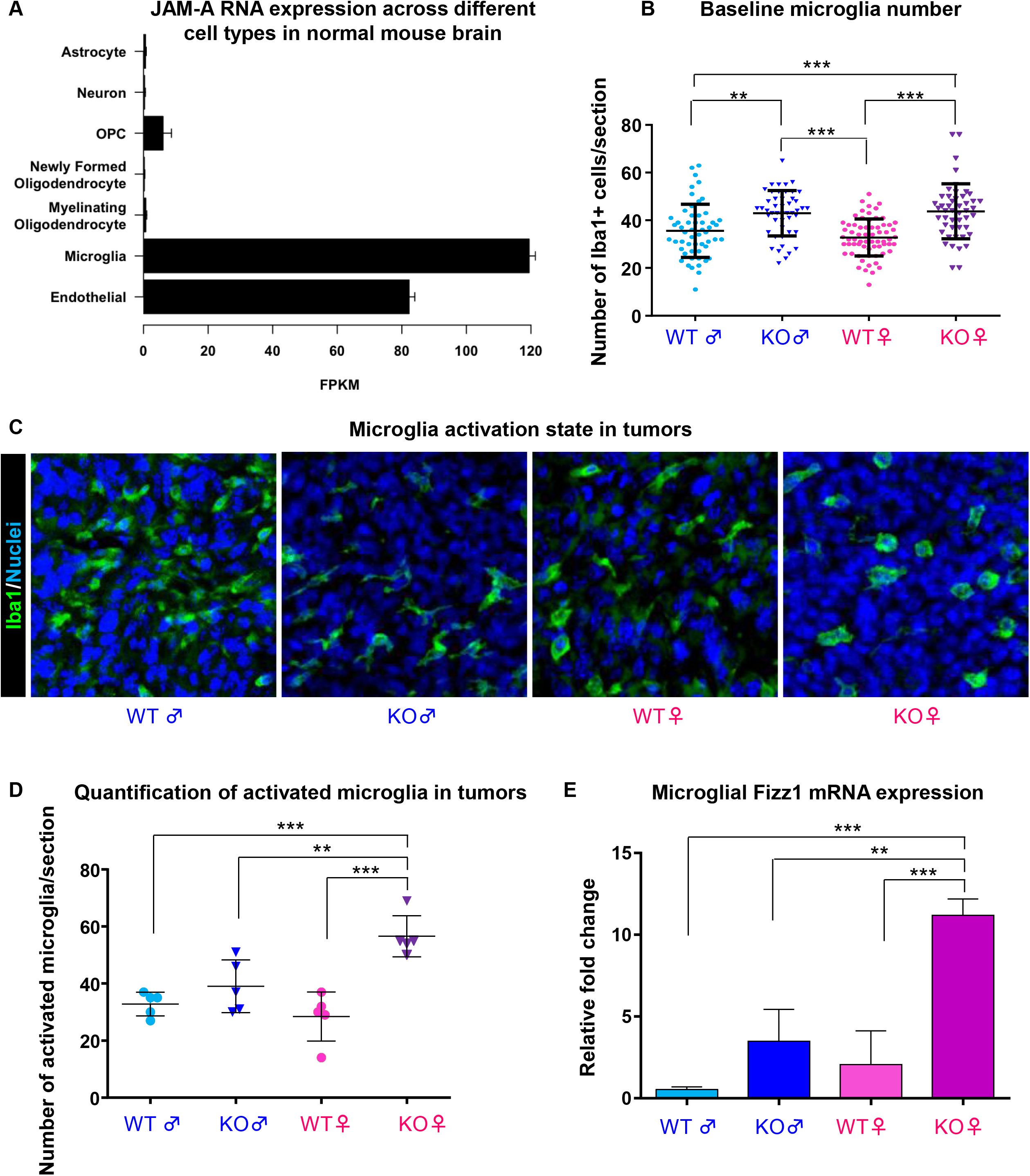
JAM-A deficiency promotes activation of female tumor associated microglia. A) The Barres Brain-RNA seq database (http://www.brainrnaseq.org/) was used to determine the mRNA expression of F11r gene (JAM-A) across different brain cell populations from healthy adult mice. B) Healthy brain tissue sections from JAM-A deficient and wild-type male and female mice (n=3 per each group) were assessed for microglia number based on immunofluorescence staining for the microglia marker Iba1. Iba1+ cells were counted using ImageJ software, and the p-value was assessed by one-way ANOVA (KO female vs KO male, *\*\*p<0.01;* KO female vs WT male, *\*\*\*p<0.001;* KO male vs WT female, *\*\*\*p<0.001;* KO Female vs WT Female, *\*\*\*p<0.001)*. C) Tumor sections of JAM-A deficient and wild-type mice were stained with the microglia marker Iba1, and amoeboid/rounded morphology was used as an indicator of microglia activation status. D) Quantification of rounded microglia in tumor sections indicating JAM-A deficient female mice have more activated microglia compared to other genotypes (n=5 tumor images per each group). Iba1+ cells were counted using ImageJ software, and the p-value was assessed by one-way ANOVA (KO female vs WT male, *\*\*\*p<0.001;* KO female vs KO male, *\*\*p<0.01;* KO female vs WT female, *\*\*\*p<0.001*). E) mRNA expression of *Fizz1* in microglia (n=4 mice per each group); p-value was assessed by one-way ANOVA (KO female vs WT male and female, ***p<0.001 and KO female vs KO male, **p<0.01).

### JAM-A deficient female microglia are phagocytic and enhance glioma cell proliferation in vitro

While these data indicate the presence of increased microglial activation in JAM-A deficient, tumor-bearing female mice, we next employed in vitro functional assessments to confirm these differences in genotype- and sex-specific microglial activation. Microglia were isolated from mixed cortical cultures from young mice, and the experimental schematic is shown (**Fig. 3A**). Using phagocytosis assays, we observed that JAM-A deficient female microglia had a significantly higher capacity to phagocytose particles compared to JAM-A deficient male microglia (**Fig. 3B**). However, this difference was not observed between sexes for wild-type microglia (**Fig. 3B**). The elevated phagocytic ability of JAM-A deficient female microglia was also recapitulated using a flow cytometry-based analysis (**Supplemental Fig. 3**). To directly assess whether microglial activation impacts GBM cell growth, we co-cultured GBM cells with microglia isolated from each genotype and found that JAM-A deficient female microglia significantly enhanced GBM cell proliferation (**Fig. 3C**). As JAM-A functions to mediate cell-cell contact, we assessed whether direct contact between microglia and GBM cells was necessary for the enhanced proliferation observed in JAM-A deficient female microglia. When GBM cells were treated with microglia conditioned media, no significant difference was observed (**Supplemental Fig. 4**), suggesting that the JAM-A deficient female microglia-mediated enhancement of GBM cell growth is contact dependent. Taken together, these data suggest that JAM-A deficient female microglia are more activated and drive GBM cell growth, which underlies the aggressive phenotype observed in vivo.

**Figure 3:**
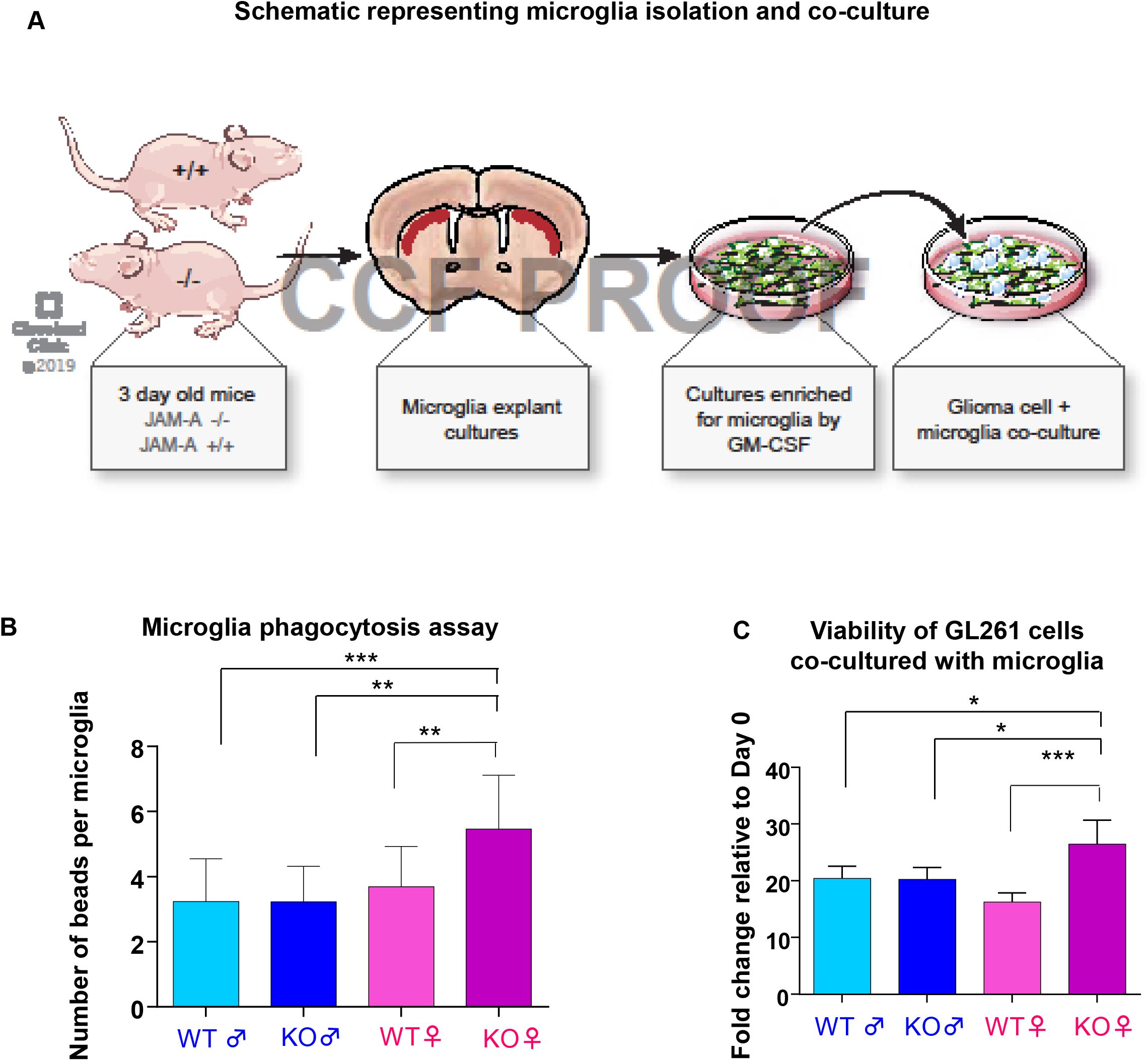
Female JAM-A deficient microglia are functionally different and drive tumor growth in vitro. A) Schematic representing microglia isolation from day 3 old JAM-A deficient and wild-type mice and co-culture with glioma cells. B) Quantitation of microglial uptake of fluorescent beads in an in vitro phagocytosis assay, using primary microglia cultures from wild-type and JAM-A deficient male and female mice. JAM-A deficient female microglia phagocytose significantly more beads compared to JAM-A deficient male and wild-type male and female microglia. No significant differences in phagocytosis were observed between wild-type male and female microglia. 20x magnification images were taken in a Keyence BZ-X fluorescent microscope and quantified using ImageJ software (15-16 images per group for WT female/male and JAM-A KO female, 9 images for JAM-A KO male). For statistical analysis, one-way ANOVA was used. Data represent mean ± SD. **p<0.01; ***p<0.001. C) Microglia from JAM-A deficient and wild-type males and females were co-cultured with GL261 cells, and proliferation of GL261 cells was measured using CellTiter-Glo proliferation assay, adjusted to a microglia alone control from each genotype. Co-culture of GL261 glioma cells with JAM-A deficient female microglia enhanced glioma cell proliferation compared to other genotypes (KO female vs WT male, *\*\*p<0.01;* KO female vs KO male, *\*\*p<0.01;* KO female vs WT female, *\*\*\*p<0.001* assessed by one-way ANOVA). n=3 mice per each group, cells plated in triplicate.

As higher levels of estrogen have been reported to increase microglial activation and lead to poorer survival in female mice with optic pathway glioma (Toonen et al., 2017b) and as estrogen has been previously reported to regulate levels of JAM-A (Choi and Baek, 2018), we wanted to test the role of estrogen in mediating the aggressive phenotype and microglial activation observed in JAM-A deficient female mice. To directly test this possibility, we assessed in vivo tumor growth in female mice after ovariectomy and observed that JAM-A deficient mice experienced a poorer survival compared to wild-type mice (**Supplemental Fig. 5A**). In fact, ovariectomy increased the aggressiveness of tumors in JAM-A deficient mice (**Supplemental Fig. 5B**). However, ovariectomy did not impact the survival of wild-type mice (**Supplemental Fig. 5C**). These data suggest that female hormones have a protective role in JAM-A deficient female mice but not in wild-type female mice.

### JAM-A deficiency in the female tumor microenvironment induces a pro-tumorigenic gene signature

To identify the mechanism through which JAM-A deficiency mediated an aggressive GBM phenotype, we employed an RNA-sequencing approach and interrogated the GL261 model transplanted into JAM-A deficient and wild-type male and female mice with the rationale that alterations in gene networks would likely be a result of cells within the tumor microenvironment, including microglia. As JAM-A deficient females had more aggressive tumors, we focused on genes that were either downregulated or upregulated compared to the other three genotypes (**Supplemental Fig. 6**). RNA-sequencing also detected an upregulation of a microglia activation marker (Retnla or Fizz1) in JAM-A deficient female tumors (**Fig. 4A**). Among the other upregulated genes, we identified an elevation in interferon activated gene 202b (Ifi202b) and validated this difference in transplanted tumors (**Fig. 4B**). Ifi202b expression was elevated in both male and female JAM-A deficient tumors compared to wild-type tumors, but we observed a significant increase in Ifi202b in JAM-A deficient female microglia compared to microglia from the other genotypes (**Fig. 4C**). To directly test whether JAM-A suppresses the expression of Ifi202b and Fizz1, we employed a function-blocking antibody to JAM-A that prevents its dimerization and subsequent downstream signaling. When JAM-A was blocked in wild-type female microglia, we observed a significant increase in Ifi202b and Fizz1 (**Fig. 4D**). These data demonstrate that JAMA deficiency in females changes the activation status of microglia via induction of Ifi202b and Fizz1 (**Fig. 4E**).

**Figure 4:**
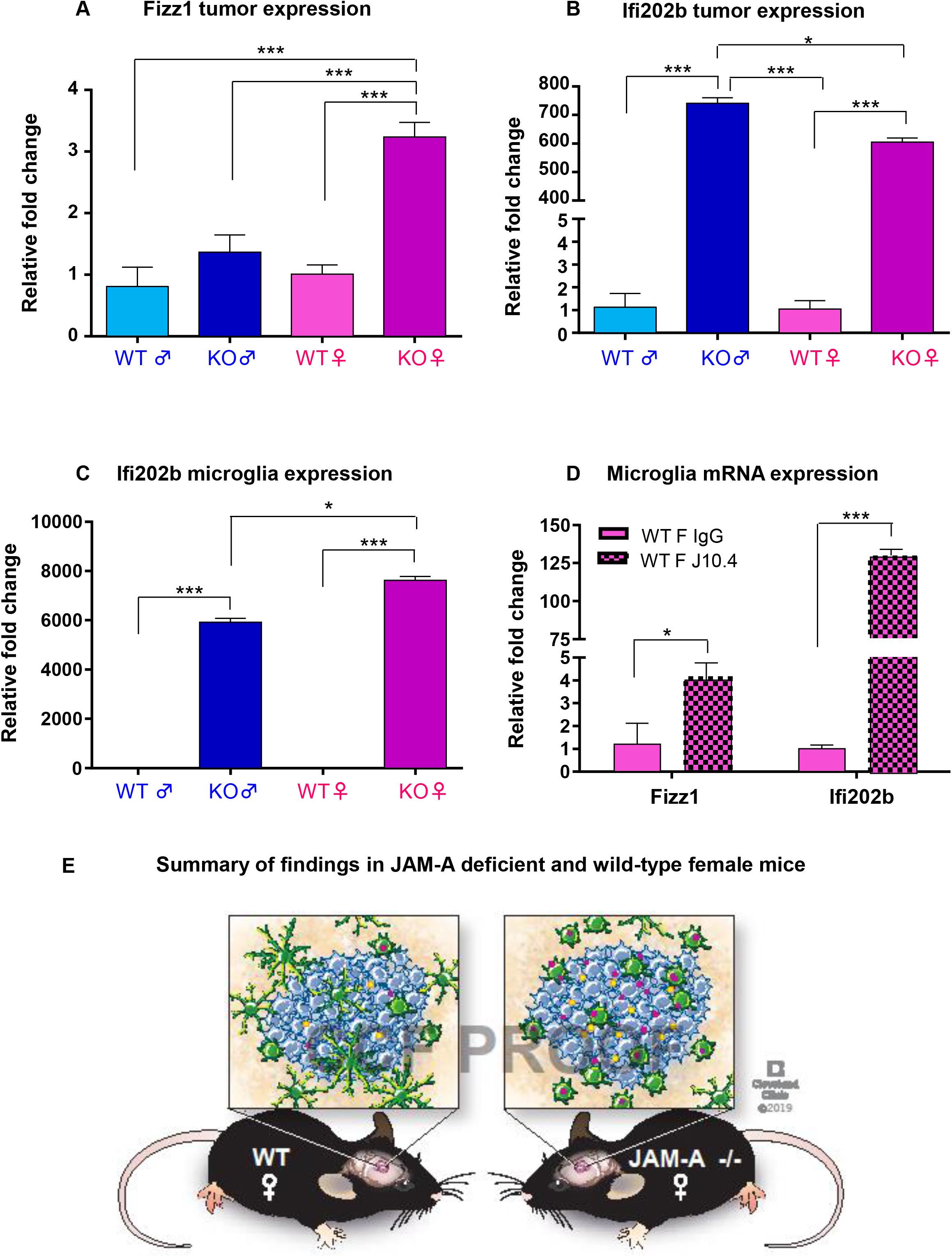
Female JAM-A deficient mice have elevated expression of Ifi202b and Fizz1 in tumor compared to female wild-type. A) *Fizz1* mRNA expression in JAM-A deficient and wild-type tumors. JAM-A deficient female tumors had higher expression compared to other groups; ***p<0.001 as assessed by one-way ANOVA. B) *Ifi202b* mRNA expression in JAM-A deficient and wild-type tumors (n=4 per each group). P-value was assessed by two-way ANOVA (KO male vs WT male and female, **p<0.001; KO male vs KO female, *p<0.05; KO female vs WT male and female, p<0.001***). C) *Ifi202b* mRNA expression in JAM-A deficient and wild-type male and female microglia (n=4 or greater per each group). JAM-A deficient female microglia had higher expression compared to all the other three genotypes. P-value was assessed by two-way ANOVA (KO female vs KO male, WT male, WT female, *\*\*\*p<0.001;* KO male vs WT male and WT female, *\*\*\*p<0.001*). D) Wild-type female microglia treated with 10 μg/ml of control IgG or J10.4 JAM-A blocking antibody. mRNA expression of *Ifi202b* and *Fizz1* was upregulated in WT female microglia (n=3 per each group) upon blocking downstream signaling of JAM-A (WT female IgG vs J10.4 *Ifi202b* expression, *\*\*\*p<0.001;* WT female IgG vs J10.4 *Fizz1* expression, *\*p<0.05*. P values were assessed by twoway ANOVA). E) Schematic representing summary of findings in JAM-A deficient and wild-type female mice. JAM-A deficient mice have more activated microglia in the tumor and express higher amounts of Ifi202b and Fizz1 compared to wild-type female mice and have poor overall survival.

While sex differences are emerging as an area of study in GBM and some key molecular alterations between male and female GBM cells have been identified, our data provide evidence that sex differences are also present in the tumor microenvironment and can impact GBM growth. This observation is consistent with sex differences observed in male and female microglia in normal and disease states (Guneykaya et al., 2018; Villa et al., 2018). Our data suggest that JAMA deficiency can induce an anti-inflammatory/pro-tumorigenic phenotype specifically in the female microenvironment, thereby increasing GBM aggressiveness. While it appears that these differences are driven more by microglia and less by TAMs or other infiltrating immune cells based on the number of cells present in the tumor microenvironment, additional functional studies assessing sex differences in immune cell interactions in GBM models is warranted. These studies could take advantage of intravital imaging approaches that have been informative in the interrogation of the dynamics of neuro-inflammation and recently adapted to GBM models (Chen et al., 2019; Davalos et al., 2008). Moreover, our findings also suggest that JAM-A functions as a tumor suppressor in the female microenvironment, while our previous work demonstrated that JAM-A is a cell-intrinsic tumor promoter in cancer stem cells (Alvarado et al., 2016; Lathia et al.,2014). These differences highlight the context-dependent role for JAM-A and provide a paradigm for the assessment of additional mechanisms that may function differently in a cell-intrinsic versus cell-extrinsic manner.

These findings also raise a series of questions that represent the starting point for future inquiry. Why does JAM-A deficiency have a stronger phenotype in females than males? In terms of activation, tumor cell proliferation, Fizz1 and Ifi202b expression, our data demonstrate that female microglia are impacted by JAM-A deficiency to a greater extent than male microglia. It is conceivable that these findings also indicate sex-specific thresholds to gene activity and function. This notion is supported by previous reports demonstrating that male astrocytes had a lower threshold for transformation than females (Kfoury et al., 2018; Sun et al., 2014). Moreover, there is strong support in the literature to suggest that immune activation status differs between males and females (Klein and Flanagan, 2016). We identified that JAM-A deficiency in female microglia enhances the expression of the anti-inflammatory genes Fizz1 and Ifi202b. Interestingly, Ifi202b was previously reported to be upregulated in mice with systemic lupus erythematosus, an auto-immune disorder to which females are more susceptible than males (Choubey and Panchanathan, 2008). IFI16, the human orthologue of Ifi202b, is involved in mediating an antiinflammatory phenotype by suppressing inflammasome activation in blood monocytes (Veeranki et al., 2011). Another outstanding question that arises from this work is the role of sex differences in cell adhesion programs. Our previous work demonstrated that adhesion is a hallmark of the stem cell state (Lathia et al., 2010; Lathia et al., 2012), but these studies were mainly focused on cell-intrinsic mechanisms and devoid of assessment between male and female GBM models. It is worth noting that the GL261 mouse model of GBM lacks sex chromosomes. Like multiple other cultured cancer cells, GL261 likely jettisoned its sex chromosomes as a result of the selective pressure of long-term culture (Xu et al., 2017). It would be interesting to revisit this hypothesis in the context of sex differences to determine whether freshly isolated male and female tumor cells have differential cell adhesion capacity. Recent work suggests that this may be the case, as longterm female GBM survivors have specific alterations in the integrin signaling network and more diffuse tumors, which may be driven by differential integrin signaling (Yang et al., 2019). As our observations demonstrate an unexpected and striking sex difference based on an adhesion mechanism in the tumor microenvironment, future studies should take into account sex differences beyond cell-intrinsic alterations. These assessment are likely to yield sex-specific data that can then be leveraged for the development of more personalized GBM therapies.

## Materials and Methods

### Animals

JAM-A knockout mice were generated by gene trap technology as previously reported (Cooke et al., 2006). Wild-type C57BL/6 male and female mice were purchased from Jackson laboratories. JAM-A (-/-) mice were bred in the animal facility and were genotyped with primer sequences previously reported (Cooke et al., 2006). All animal experiments were performed under Cleveland Clinic-approved Institutional Animal Care and Use Committee (IACUC) protocols.

### Intracranial implantation of tumor cells

Six-week-old male and female JAM-A deficient and wild-type male and female mice were used for all experiments. A total of 10,000 GL261 or 5,000 SB28 cells were resuspended in 5 μl of RPMI null media and injected intracranially into the left hemisphere 2 mm caudal to the coronal suture, 3 mm lateral to the sagittal suture at a 90° angle with the murine skull to a depth of 2.5 mm. Mice were monitored daily for signs of tumor burden and sacrificed when symptomatic.

### Immunofluorescence

Immunostaining analysis of GL261 tumor specimens (generated by intracranial injection) was performed as following. Transplanted mouse brains containing tumors were dissected from JAM-A deficient and wild-type mice, fixed in 4% formaldehyde, dehydrated using 30% sucrose, and embedded in optimum cutting temperature medium (OCT, Tissue-Tek). Ten micrometer tissue sections harboring tumors were mounted on glass slides (Superfrost microscope slides, Fisher Scientific), and cells were permeabilized using blocking buffer (PBS with 10% normal goat serum and 0.1% Triton X-100). Cells were washed, and primary (Anti-Iba1 (1:100) Wako chemicals (019-19741)) and secondary antibodies (Alexa Flour 488 (1:500)) were diluted in blocking buffer prior to staining. Cell nuclei were stained with Hoechst 33342 at a 1:10,000 dilution in PBS, coverslips were mounted using Gelvatol Mounting Reagent, and slides were stored at −20°C until imaging. Slides were imaged using the 45x lens of Keyence BZ-X fluorescent microscope microscope. All images are representative of four random fields within the stained sample and representative of three separate experiments. Microglia counting was performed using Image J, rounded or amoeboid microglia with no ramifications were considered as activated.

### Primary microglia cultures

Mixed cortical cultures from newborn JAM-A deficient and wild-type mice (day 0–3) were generated using standard protocols (Marshall et al., 2008). Neurogenic astrocytic monolayer cultures of cells at passage 1-3 were grown to confluency, after which the neural growth media was replaced with microglial proliferation media (MPM) consisting of DMEM/F12, 10% FBS, N2 supplement, and recombinant mouse granulocyte macrophage-colony stimulating factor (GM-CSF) at a concentration of 20 ng/mL (murine rGM-CSF, Peprotech 315-03). MPM was replaced every three days until phase-bright microglia became visible in the adherent culture and floater cells became visible in the media. Propagated cultures were agitated at room temperature at 100 rpm for 30 minutes, after which the media and the detached cells were collected for analysis.

### Microglia phagocytosis assay

Microglia phagocytosis assays were performed according to previously published protocols (Lian et al., 2016). Microglia (50,000 cells) isolated from JAM-A deficient and wild-type male and female mice were plated onto a coverslip, incubated with GFP-labelled fluorescent beads (L1030,Sigma Aldrich) and fixed, and microglia were stained with anti-Iba1 antibody. Imaging was done using Keyence BZ-X fluorescent microscope and quantification of microglia uptake of beads was done using ImageJ software.

### Microglia conditioned media-glioma cell proliferation assay

Microglia were plated on a 6-well plate and incubated for 48 hr, after which conditioned media was collected. GL261 cells were resuspended in microglia conditioned media, and 1000 cells/well were plated in a 96-well plate in triplicate. Proliferation of cells was measured using CellTiter-Glo (Promega) on days 0, 1, 3 and 7 according to the manufacturer’s protocol, which uses ATP content as a surrogate of cell number.

### Microglia-glioma co-culture proliferation assay

For co-culture cell proliferation assay, 1,000 GL261 and 2500 microglia cells were plated per well in RPMI growth media in a white-walled 96-well plate in triplicate. The number of cells was measured using CellTiter-Glo (Promega) on days 0, 1, 3 and 7. As controls, we individually assessed microglia proliferation alone, and this proliferation fold change was subtracted from the respective co-cultures of each group.

### RNA-sequencing analysis

RNA was extracted using an RNeasy mini kit from JAM-A deficient and wild-type male and female tumors (n=3 per each group). Total RNA (50 ng) was used to generate whole transcriptome libraries for RNA sequencing using Illumina’s TruSeq RNA Sample Prep. Poly(A) mRNA selection was performed using oligo(dT) magnetic beads, and libraries were enriched using the TruSeq PCR Master Mix and primer cocktail. Amplified products were cleaned and quantified using the Agilent Bioanalyzer and Invitrogen Qubit. The clustered flowcell was sequenced on the Illumina HiSeq 4000 for paired 100-bp reads using Illumina’s TruSeq SBS Kit V3. Lane level fastq files were appended together if they were sequenced across multiple lanes. These fastq files were then aligned with STAR 2.4.0 to the mm10 mouse reference genome. Transcript abundance was quantified and normalized using Salmon in the unit of transcripts per million (TPM). Consensus clustering was performed to assess overall similarity among samples using the ConsensusClusterPlus R package, and heatmaps were generated using the R heatmap.2 package with Euclidean Distance and average clustering method. Gene set variation analysis (GSVA) (Hanzelmann et al., 2013) was then performed to determine the change and variation in pathway activities of JAM-A deficient female mice. Log transformation was applied, and default parameter settings were followed for RNA-seq data in GSVA.

### qRT-PCR

RNA from cells of interest was extracted using an RNeasy mini kit, and cDNA was synthesized using qSCRIPT cDNA Super-mix (Quanta Biosciences). qPCR reactions were performed using an ABI 7900HT system using Fast SYBR-Green Mastermix (SA Biosciences, Valencia, CA, USA). For qPCR analysis, the threshold cycle (CT) values for each gene were normalized to the expression levels of CycloA. Dissociation curves were evaluated for primer fidelity, and only threshold cycles below 35 cycles were reported. All qRT-PCR experiments were performed at least three times. The following primers (Integrated DNA Technologies) were used:

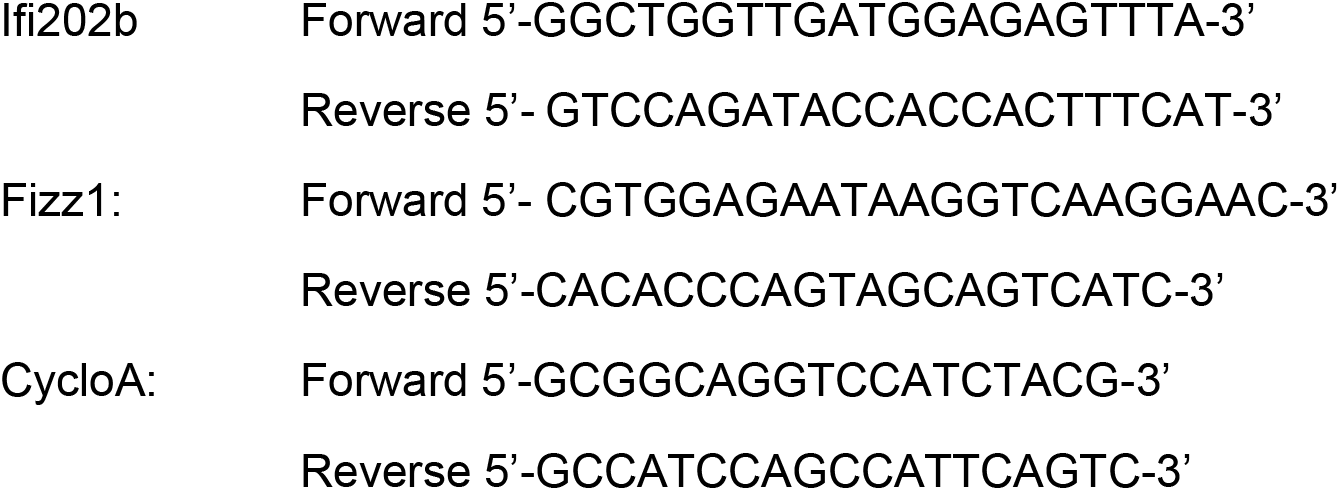

### Sex-determination PCR

For sex-determination of mice, tails from 1-3-day-old JAM-A deficient and WT mice were collected, and DNA was isolated. Sex of microglia was confirmed by expression of the X- and Y-encoded paralogs Jarid1c and Jarid1d using previously published protocols (Clapcote and Roder, 2005).

### Ovariectomy

Female mice at the age of 4-5 weeks were used for ovariectomy. Ovariectomies were performed via the dorsal route. The skin incision was pulled approximately 0.5 cm horizontally to the left or the right to identify two fat masses next to the lower pole of the kidneys. The left and right ovaries of the mouse are found in the respective fat masses, and they were bilaterally ovariectomized (Yan et al., 2018). For the sham control group, the dorsal skin and abdominal wall were incised, and the ovaries were directly visualized but not removed. Animals were allowed to recover for 2 weeks and were then used for intracranial tumor injections.

### JAM-A blocking antibody treatment of microglia

A total of 3-4 x 10^5^ microglia were plated in a 6-well plate and treated with 10 μg/ml immunoglobulin IgG1 control antibody (Santa Cruz Biotechnology, sc-2025) or JAM-A-blocking antibody J10.4 (Santa Cruz Biotechnology, sc-53623). Cells were incubated for 48 hr, mRNA was isolated using an RNeasy mini kit, and qPCR was performed.

### Flow cytometry analysis

Aged-matched JAM-A^-/-^ and WT female mice were implanted with GL261 tumors as described above. Sixteen days post-tumor implantation, mice were euthanized, and blood was collected into Safe-T-Fill Capillary Blood Collection Tubes (DevineMedical) via cardiac puncture. Single-cell suspensions from tumor-bearing and contralateral hemispheres were prepared via mechanical dissociation. Samples were strained through 40 μm filters (Fisher scientific) and washed with PBS twice. Cells were first stained with LIVE/DEAD Fixable Dead Cell Stain Kits (ThermoFisher Scientific) based on manufacturer’s instructions and then incubated with FcR Blocking Reagent (Miltenyi Biotec) diluted in 2% FBS in PBS. Cells were incubated on ice with antibody cocktails containing Ly6C, Ly6G, CD11b, CD11c, I-A/I-E, CD68, CD45, CD4, CD3 and CD8 (Biolegend) for 20 minutes on ice. Samples were fixed with eBioscience™ Foxp3/ Transcription Factor Staining Buffer Set (Thermo Fisher Scietific) and were acquired via a BD LSRFortessa (BD Biosciences). FlowJo software was used for sample analysis (BD Life Sciences).

### Statistical Analyses

Graphs were created using GraphPad Prism 6.0. Results are expressed as mean ± SD. Information regarding the numbers of experimental replicates, statistical tests performed, and significance values can be found in the figure legend for each figure panel. *\*p < 0.05, **p <0.01, ***p < 0.001* indicates statistical significance.

## Supporting information

Supplementary data revised

## Acknowledgements

We thank the members of the Lathia and Davalos laboratories for insightful discussion and constructive comments on the manuscript. We thank the Thesis Committee members of ST: Drs. Crystal Weyman, Thomas McIntyre, Alexandru Almasan and Hannelore Heemers for their valuable and constructive comments on the manuscript. We thank Amanda Mendelsohn and the Center for Medical Art and Photography at the Cleveland Clinic for providing illustrations and Dr. Erin Mulkearns-Hubert for editorial assistance. This work was funded by the NIH (R01 NS083629 (JDL) and R01 NS112526 (DD)), Sontag Foundation (JDL), Cleveland Clinic Brain Tumor Center of Excellence (JDL), Cleveland Clinic VeloSano Bike Race (DD, JDL), Case Comprehensive Cancer Center (JSB-S, JDL), American Brain Tumor Association Research Collaboration Grant (JBR, JDL), Cleveland State University Graduate Student Research Award (ST), and Case Comprehensive Cancer Center T32 CA059366-23 (DB),1F32CA213727-01 (DJS) and the Bodossaki Foundation (EP).

