## Supplementary data revised for "JAM-A functions as a female microglial tumor suppressor in glioblastoma"

#### S Fig1 Survival data upon intracranial injection of tumor cells

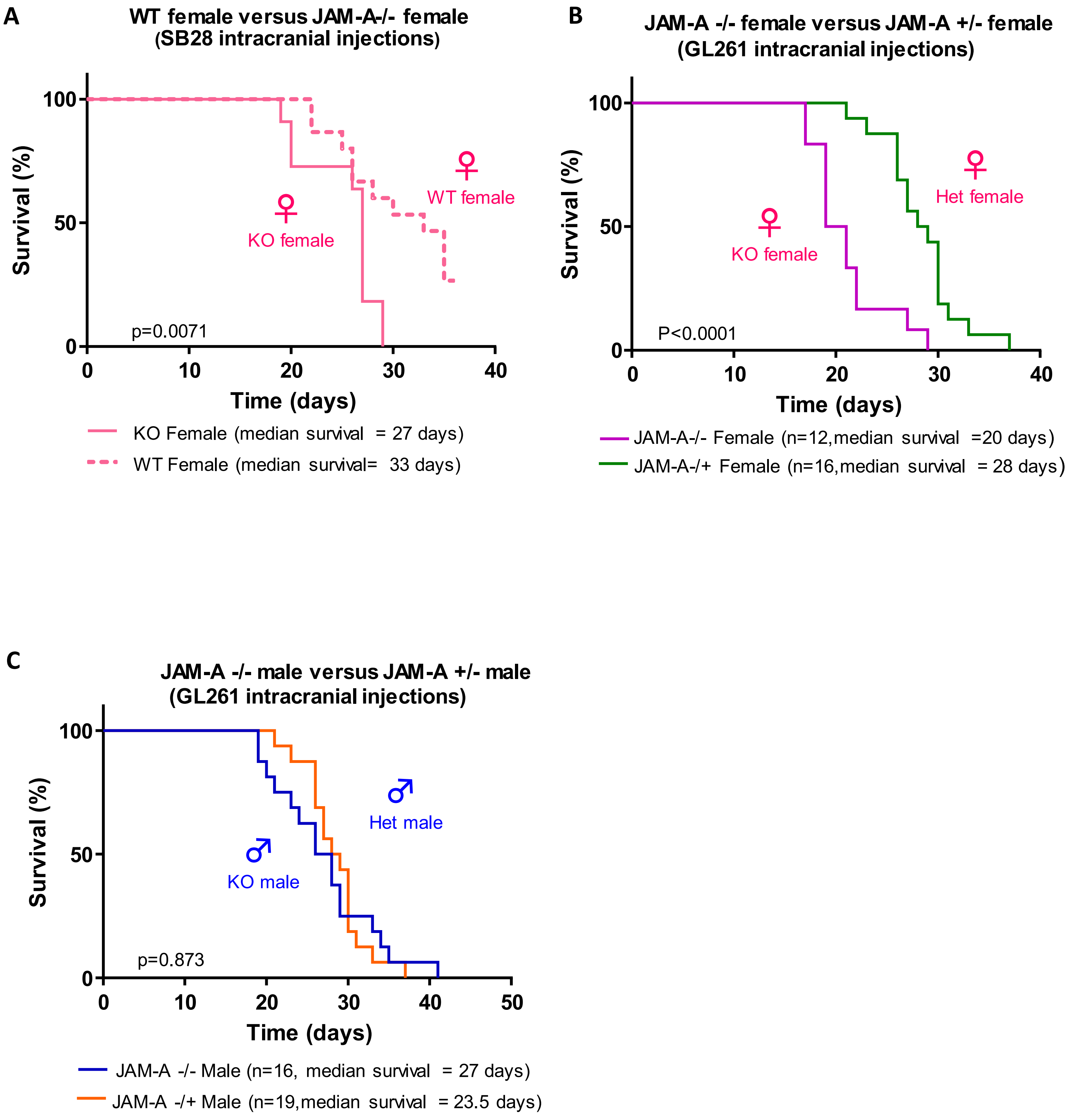

#### S Fig2 Immune cell profiling in JAM-A deficient and wild-type females

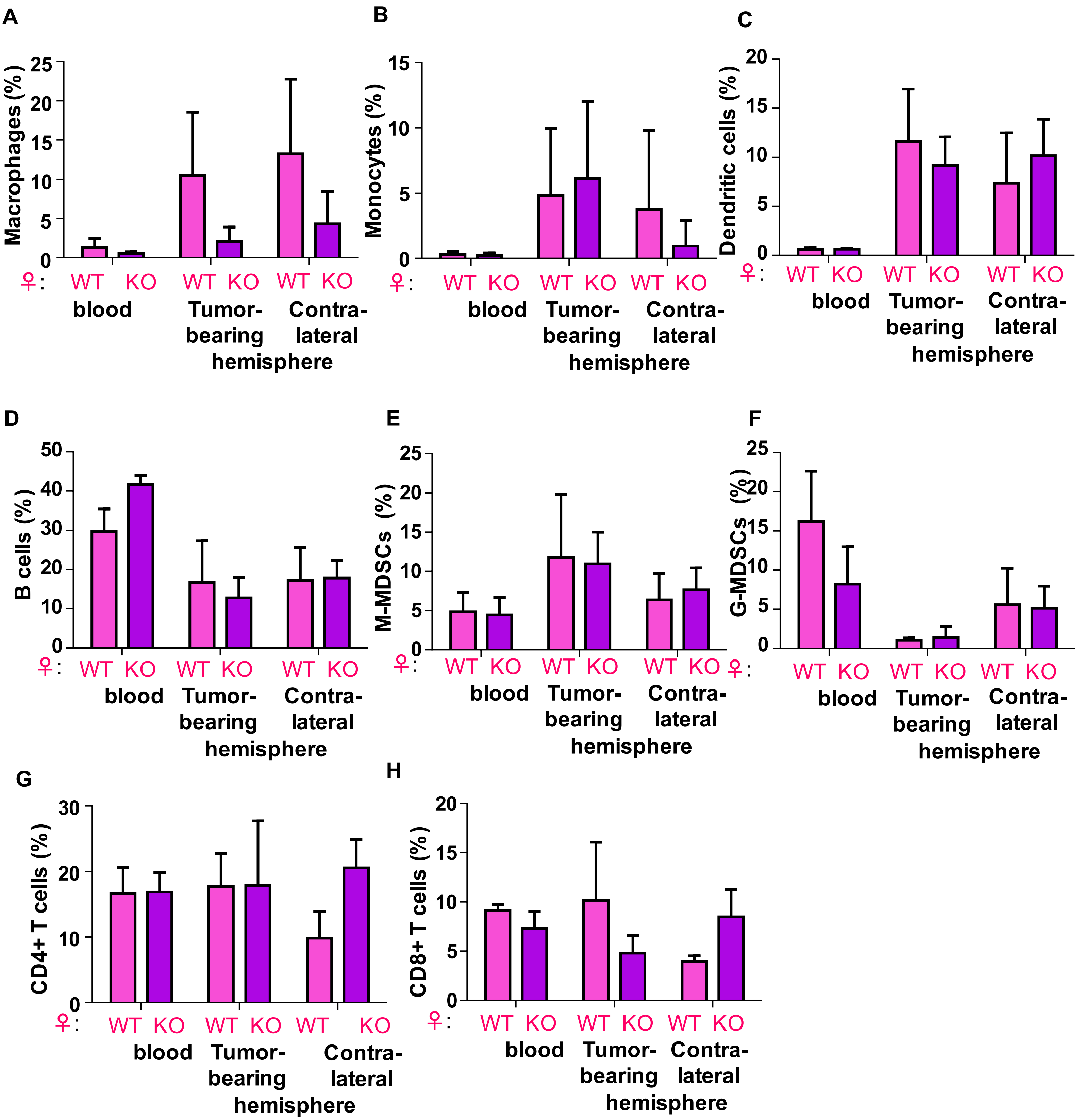

### S Fig3 Microglia phagocytosis assay by flow cytometry

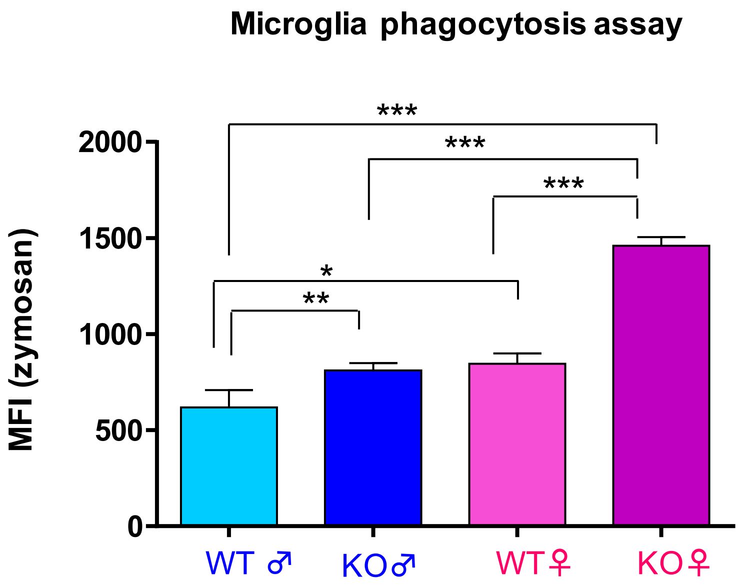

#### S Fig4 Microglia and GL261 microglia conditioned media proliferation assay

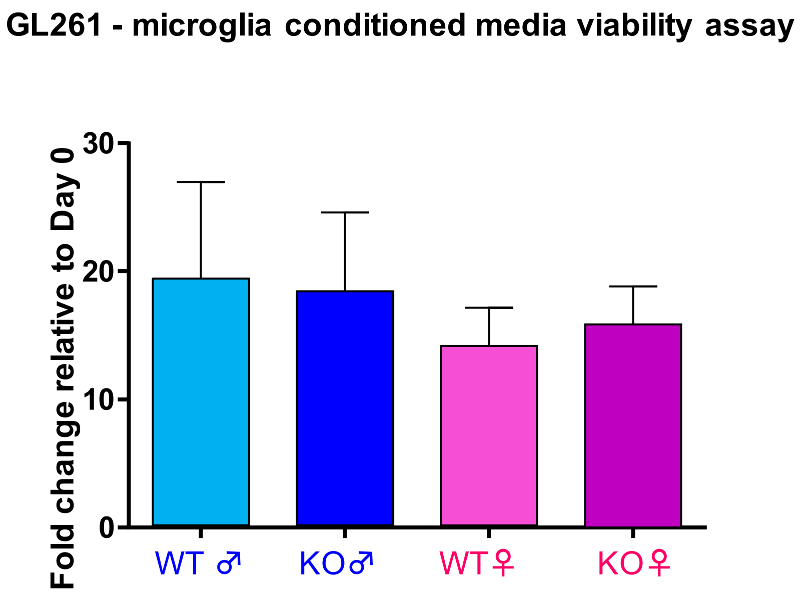

**S Fig5 Survival upon ovariectomy and intracranial implantation of tumor cells**

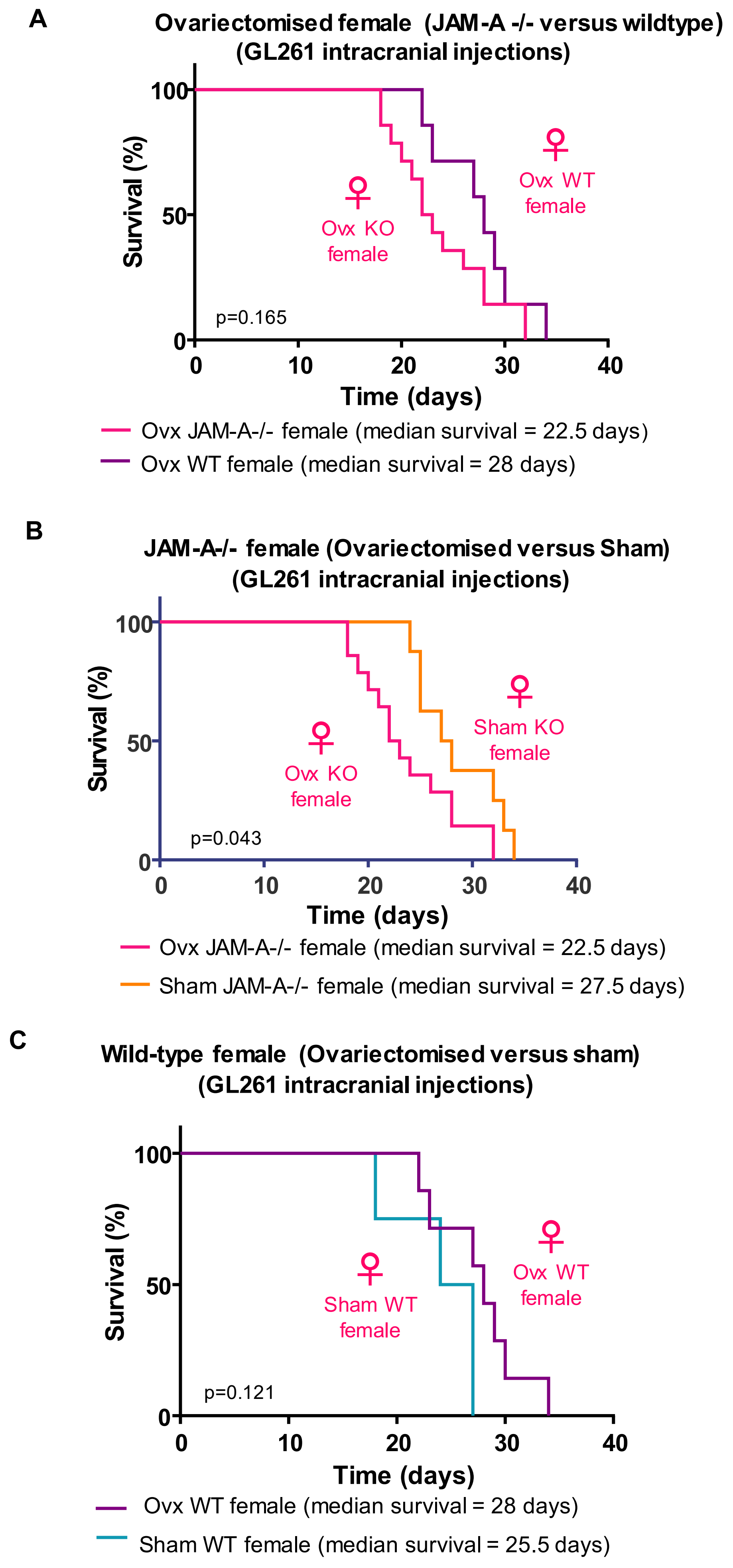

#### S Fig6 RNA Sequencing reveals gene programs that are unique to tumors in JAM -A deficient mice

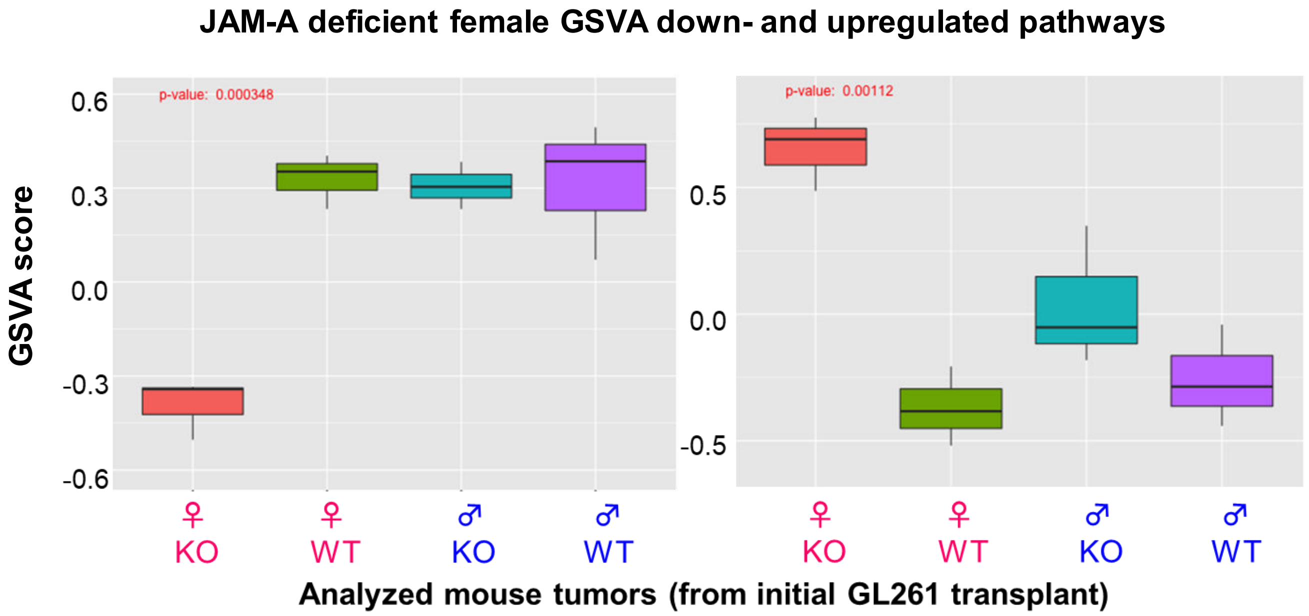

**Supplementary Figure Legends**

**Supplementary Figure 1: Survival curves upon intracranial injection of tumor cells.**
A) Survival curves upon intracranial implantation of SB28 glioma cells into JAM-A deficient female (n=16, median survival=27 days) and wild-type female mice (n=10, median survival=30 days); **p=0.024.* B) Survival curves upon intracranial implantation of GL261 glioma cells into JAM-A deficient female (n=12, median survival=20 days) and heterozygous female mice (n=16, median survival=28 days), ****p<0.001*. C) Survival curves upon intracranial implantation of GL261 glioma cells into JAM-A deficient male (n=16, median survival=27 days) and heterozygous male mice (n=19, median survival=23.5 days); *p=0.873*. Kaplan-Meier survival curves were plotted using GraphPad Prism 6.0, and p-values were assessed using log-rank (Mantel-Cox) test.

**Supplementary Figure 2: Immune cell profiling of JAM-A deficient and wild-type females.**
Percentage of A) macrophages, B) Monocytes, C) Dendritic cells, D) B cells, E) M-MDSCs, F) G-MDSCs, G) CD4+ T cells, and H) CD8+ T cells in wild-type and JAM-A deficient female mice blood and brain (tumor and contralateral hemisphere). p-values were assessed by two-way ANOVA; no significant differences were observed across other immune cell types between the two groups.

**Supplementary Figure 3: Microglia phagocytosis assay by flow cytometry**
Microglial cells (50,000 cells) from JAM-A deficient and wild-type male and female mice were treated with pHrodo green zymosan bioparticles (30 µg/ml) (ThermoFisher # P35365) and incubated for 30 mins, after which their uptake was assessed by flow cytometry. Samples were analyzed with a BD Fortessa, and zymosan uptake was quantified based on changes in the mean fluorescence intensity using FlowJo software. p-values were assessed by one-way ANOVA (KO female vs KO male and WT male and female, ***p>0.001, WT female vs WT male, *p<0.05, WT female vs KO male, **p<0.01).

**Supplementary Figure 4: Microglia and GL261-microglia conditioned media proliferation assay**
GL261 cells (1000 cells) were treated with microglia conditioned media, and cell proliferation was measured at Day 7 using the CellTiter-Glo proliferation assay. The p-values were measured by one-way ANOVA, and no significant differences in proliferation were observed.

**Supplementary Figure 5: Survival upon ovariectomy and intracranial implantation of tumor cells**
A) Survival curves upon ovariectomy and intracranial implantation of GL261 cells in JAM-A deficient (n=14, median survival=22.5 days) and wild-type female mice (n=7, median survival=28 days); p=0.165. B) Survival curves between ovariectomized JAM-A deficient females (n=14, median survival=22.5 days) and sham JAM-A deficient females (n=8, median survival=27.5 days); *p=0.043. C) Survival curves between ovariectomized wild-type female (n=7, median survival=28 days) and sham wild-type female mice (n=4, median survival=27.5 days); p=0.121. Kaplan-Meier survival curves were plotted using GraphPad Prism 6.0, and p-values were assessed using log-rank (Mantel-Cox) test.

**Supplementary Figure 6: RNA Sequencing reveals gene programs that are unique to tumors in JAM-A deficient mice**

RNA-Sequencing of GL261 tumors from JAM-A deficient and wild-type male and female mice (n=3 mice per each group). Genes specifically up- and downregulated in JAM-A deficient tumors are represented according to their GSVA score.
